# A Cis-Element in *magu-2* Mediates Apical mRNA Localization via Active Transport in *C. elegans*

**DOI:** 10.64898/2026.10.17.759870

**Authors:** Zhimin Li, Qinghao Jiang, Zirui Wang, Yu Sun

## Abstract

mRNA subcellular localization is a key mechanism of post-transcriptional gene regulation and is widely conserved across cell types and species. Although increasing evidence suggests that transcripts associated with epithelial polarity also undergo asymmetric subcellular distribution, the trafficking patterns and regulatory mechanisms governing this process remain elusive. MAGU-2, the *Caenorhabditis elegans* ortholog of the *Drosophila* polarity gene stardust (*sdt*), has previously been reported to demonstrate apical mRNA enrichment. In this study, a truncation screen of *magu-2* exon 3 was performed, leading to the identification of a 270-nt SH3 domain-containing fragment that drives apical localization of both its encoded mRNA and the corresponding truncated MAGU-2 protein, indicating that this 270-nt fragment within magu-2 is sufficient to direct the apical localization of both the transcript and its protein product. Live imaging, RNAi-mediated knockdown of DHC-1, BicD, or EGAL-1, together with nocodazole treatment, revealed that this mRNA undergoes active transport and that its apical targeting is disrupted upon impairment of dynein/BicD/Egalitarian function or microtubule integrity is compromised, implicating the conserved dynein/BicD/Egalitarian axis in *magu-2* mRNA trafficking. While deletion of this 270-nt element exerted no overt effect on epithelial polarity or development, these findings provide insight into the active transport of epithelial polarity-related transcripts and position *magu-2* as a useful model for dissecting cis-elements and trans-acting factors modulating subcellular RNA localization.

## 1. Introduction

mRNA localization is the asymmetric distribution of transcripts within the cytoplasm following nuclear export. This evolutionarily conserved process is widespread across diverse cell types and species, playing a pivotal role in post-transcriptional gene expression regulation [1]. Genome-wide analysis of mRNA localization in *Drosophila* embryos identified subcellular localization for 71% of 3,370 transcripts, uncovering diverse and specific mRNA localization patterns [2]. At the cellular level, mRNA subcellular localization plays essential roles in regulating cell polarity, differentiation, and migration. For instance, the apical localization of *ZO-1* mRNA mediates epithelial tight junctions in mammals [3]. Beyond individual cells, this process exerts profound effects on organ and organismal homeostasis, tissue differentiation, and development [4-7]. Notably, mRNA mislocalization can precipitate pathological conditions, including neurodegenerative diseases, muscular dystrophy, and cancer [6, 8, 9].

Studies in model organisms have revealed three canonical modes of RNA transport: (i) passive diffusion and anchoring, wherein transcripts explore the cytoplasm by Brownian motion before cytoskeletal capture. While this model is universal, it is kinetically inefficient and insufficient to establish robust asymmetry; (ii) active transport, in which localization elements bearing distinct secondary structures recruit RNA-binding proteins (RBPs) to assemble ribonucleoprotein (RNP) complexes, which are shuttled either by canonical motor proteins (e.g., kinesin, dynein) or by motile organelles (e.g., endosomes, lysosomes), to drive long-range directed movement along cytoskeletal tracks. For example, in *Drosophila* oocytes, microtubule-based transport of *oskar* mRNA relies on the adaptor proteins Egalitarian and Bicaudal-D to load the transcript onto dynein motors [10-13]. Recently, TIA1-containing RNPs have been described to undergo retrograde axonal transport by dynein, mediated by the calcium-sensitive adaptor ANXA7 [14]; (iii) selective degradation, whereby localized degradation enhances spatial specificity(Chen et al., 2014).Thus, identification of the cis-acting elements and transport complexes that direct mRNA trafficking, coupled with dynamic tracking of transcript translocation, is essential for elucidating the mechanisms underlying subcellular localization.

In recent years, accumulating evidence has revealed that transcripts associated with epithelial polarity exhibit mRNA subcellular localization [15]. In the intestinal epithelium, approximately 30% of highly expressed transcripts exhibit an apical-basal polarized distribution [16]. Meanwhile, in both *Drosophila* and human cell lines, mRNAs encoding polarity determinants such as ZO-1, Afdn, and Scrib are localized to cell junction regions [17]. Similarly, in the epithelial cells of *Caenorhabditis elegans*, multiple membrane-associated mRNAs are enriched at the cell cortex during early embryonic development, and among transcripts encoding proteins related to the *C. elegans* apical junction (CeAJ), *dlg-1, ajm-1, erm-1*, and *sma-1* mRNAs have been reported to display subcellular localization [18, 19]. These findings suggest that epithelial polarity is achieved not only through traditional protein localization but also partly through mRNA transport and local translation, constituting an alternative route for protein targeting.

Epithelial polarity manifests as an intrinsic apicobasal asymmetry that partitions cells into apical and basolateral domains, characterized by the asymmetric distribution of proteins and lipids along this axis. At the apical-lateral interface, cell-cell junctions sustain cell cohesion, scaffold polarity regulators, and demarcate the membrane boundaries [20]. Apicobasal polarity maintains epithelial homeostasis, while its dysregulation contributes to developmental disorders and carcinoma progression [21].

A multilayered antagonistic network between conserved polarity modules specifies epithelial apicobasal polarity and defines membrane domain identities [22]. Apical polarity is established and maintained by the PAR complex (PAR-3/PAR-6/aPKC), which integrates Cdc42 signaling to phosphorylate and exclude basolateral determinants, and by the Crumbs complex, which defines the apical membrane domain and provides a scaffold for the cortical actin cytoskeleton [23].

In vertebrates, the Crumbs complex comprises the transmembrane receptor Crumbs, the scaffolding protein PALS1, and the adaptor proteins PATJ/LIN7 [24]. PALS1 contains four conserved domains, namely L27, PDZ, SH3, and GUK, among which SH3 and PDZ domains, directly engages the C-terminal ERLI motif of Crb3, with this interaction is further stabilized by the GUK domain. In contrast, the N-terminal L27 domain recruits the adaptor proteins PATJ and LIN7 to the apical–lateral marginal zone, thereby anchoring the Crumbs complex in this domain [25, 26]. Notably, PALS1 is essential for apical domain homeostasis, tight junction anchoring, and tissue morphogenesis in *Drosophila*, zebrafish, mice, and humans. Its loss or mislocalization of which dismantles the Crumbs complex and drives tissue-specific pathologies, including mixed CRB1/CRB2-type retinal degeneration, renal cyst formation, and neural tube defects [27-29]. The *C. elegans* homolog of PALS has been identified as MAGU-2, which contains the conserved PDZ, SH3, and GUK domains[30]. A previous work demonstrated that exon 3 of *magu-2*, which encodes the PDZ and SH3 domains, is both necessary and sufficient to drive *magu-2* mRNA and protein to the apical membrane of intestinal cells in *C. elegans* [31].

In this study, we investigated the regulation and biological significance of *magu-2* mRNA localization in *C. elegans* intestinal cells. Truncation screening of *magu-2* exon3 mapped its apical targeting signal to a 270-nt fragment containing the SH3-coding region, which harbors a localization element that induces both mRNA and protein co-localization to the intestinal apical domain. Live-cell imaging further unveiled that this transcript exhibits directed movement, indicative of active RNA transport. Furthermore, RNAi-mediated knockdown of *dhc-1, bicd* or *egal*, as well as drug-induced microtubule defects, disrupted the apical targeting of 270-nt fragment-derived *magu-2* mRNA and protein, yielding discrete cytoplasmic aggregates and diffusion. However, deletion of this 270-nt fragment in *C. elegans* produced no overt defects in intestinal epithelial cells, nor detectable alterations to developmental abnormalities.

## 2. Results

### 2.1. Two independent sequences in magu-2 exon 3 are themselves sufficient for apical localization

In *C. elegans*, a previous study has demonstrated that exon 3 of *magu-2*, which encodes the PDZ and SH3 domains, yields both transcripts and the encoded protein enriched at the apical membrane [31]. To define the sequence determinants underlying apical localization, a series of magu-2 exon 3 truncations was designed based on its PDZ1-SH3 domain architecture. Each reporter encoded GFP, harbored one of the truncated exon 3 fragments, included a 24 × PP7 stem-loop array for live-cell mRNA labeling, and terminated with the unc-54 3’UTR; multiple independent transgenic lines were established by microinjection (Fig. 1A). Subcellular localization analyses revealed that the protein encoded by full-length exon 3 was enriched at the apical membrane, consistent with the finding of previous report (Fig1.B) [31]. Notably, overexpression of the PDZ-SH3 region induced dramatic intestinal morphological defects characterized by expansion of the apical domain — a phenotype reminiscent of that observed upon overexpression of *crb-3* or *eat-20* (Fig.1C) [30]. Although the PDZ domain when expressed alone failed to target to the apical membrane, the polypeptide encoded by a 270-nt sequence spanning from the region immediate downstream of the PDZ domain to the SH3 domain was efficiently localized to the apical membrane, consistent with the distribution pattern of full-length exon 3(Fig.1G-H, Fig.1SA). Further truncation of this 270-nt fragment revealed that neither the 5’ half nor the 3’ half (each 135 nt) was sufficient for apical targeting; both truncated fragments formed punctate structures dispersed throughout the cytoplasm, precluding further systematic dissection (Fig.1I, J). Interestingly, deletion of the entire PDZ-SH3 module did not abolish apical localization (Fig. 1D). Subsequent fine-scale mapping identified a 354-nt sequence immediately upstream of the PDZ domain as sufficient to drive apical enrichment (Fig. 1E, Fig.1SB).

**Figure 1.**
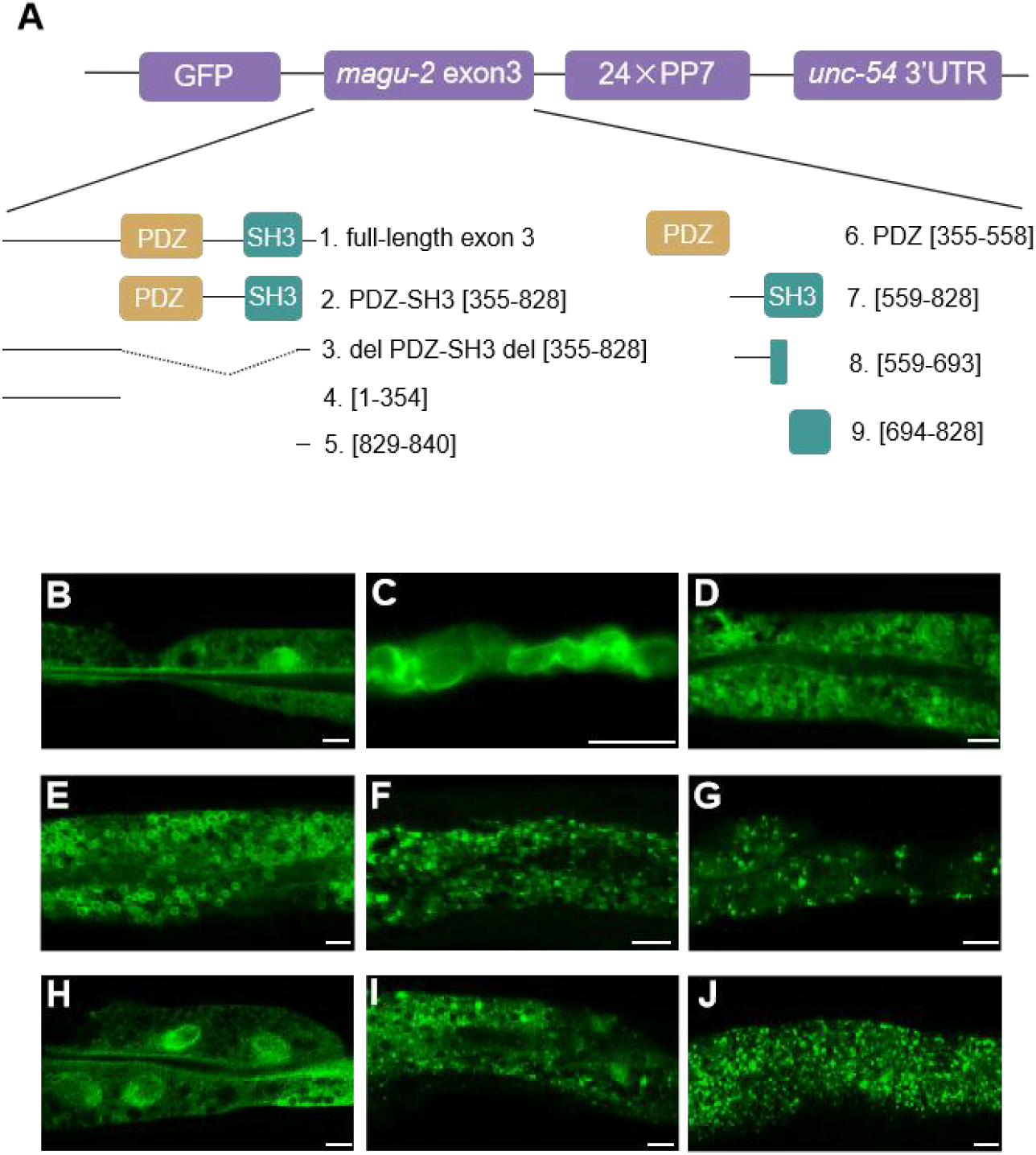
magu-2 exon 3 contains two independent apical localization elements. (A)Schematic of nine GFP-tagged magu-2 exon 3 truncation constructs containing the 24×PP7 cassette and unc-54 3’UTR. (B-J) Localization of proteins expressed by GFP-tagged constructs 1-9 in C. elegans intestinal cells. (B) Full-length exon 3 shows apical enrichment of the GFP-tagged protein. (C) Overexpression of the PDZ-SH3 construct causes intestinal tube defects. (D) Exon 3 lacking the PDZ-SH3 domain remains apically enriched. (E) The N-terminal region of exon 3 upstream of PDZ exhibits apical enrichment. (F) The region downstream of the SH3 domain in exon 3 shows the protein distributed as puncta in the cytoplasm in a disordered pattern (G) The PDZ domain displays punctate, disordered cytoplasmic distribution. (H) The 270-nt region downstream of PDZ, containing the SH3 domain, exhibits a distribution consistent with exon 3. (I-J) The 5’and 3’ halves of the 270-nt region downstream of PDZ both show punctate, disordered cytoplasmic distribution. Scale bar: 5 μm for all images

### 2.2. A 270-nt core fragment of magu-2 exon 3 is sufficient for mRNA localization to the apical membrane in intestinal epithelia

To map the mRNA localization patterns of *magu-2* exon 3 truncations, and building on extend prior observations of protein localization phenotypes, single-molecule fluorescence in situ hybridization (smFISH) was conducted. Briefly, smFISH probes complementary to the GFP coding sequence were designed to enabled unbiased detection of all truncation-derived transcripts, as GFP is present in every reporter construct. The panel of reporters tested included: (i) the region upstream of the PDZ domain; (ii) the PDZ coding sequence; (iii) the 270-nucleotide (nt) core fragment spanning the PDZ C-terminus to the SH3 domain; and (iv) 5’- and 3’-135 nt truncations of this 270-nt fragment (Fig. 2A).

**Figure 2.**
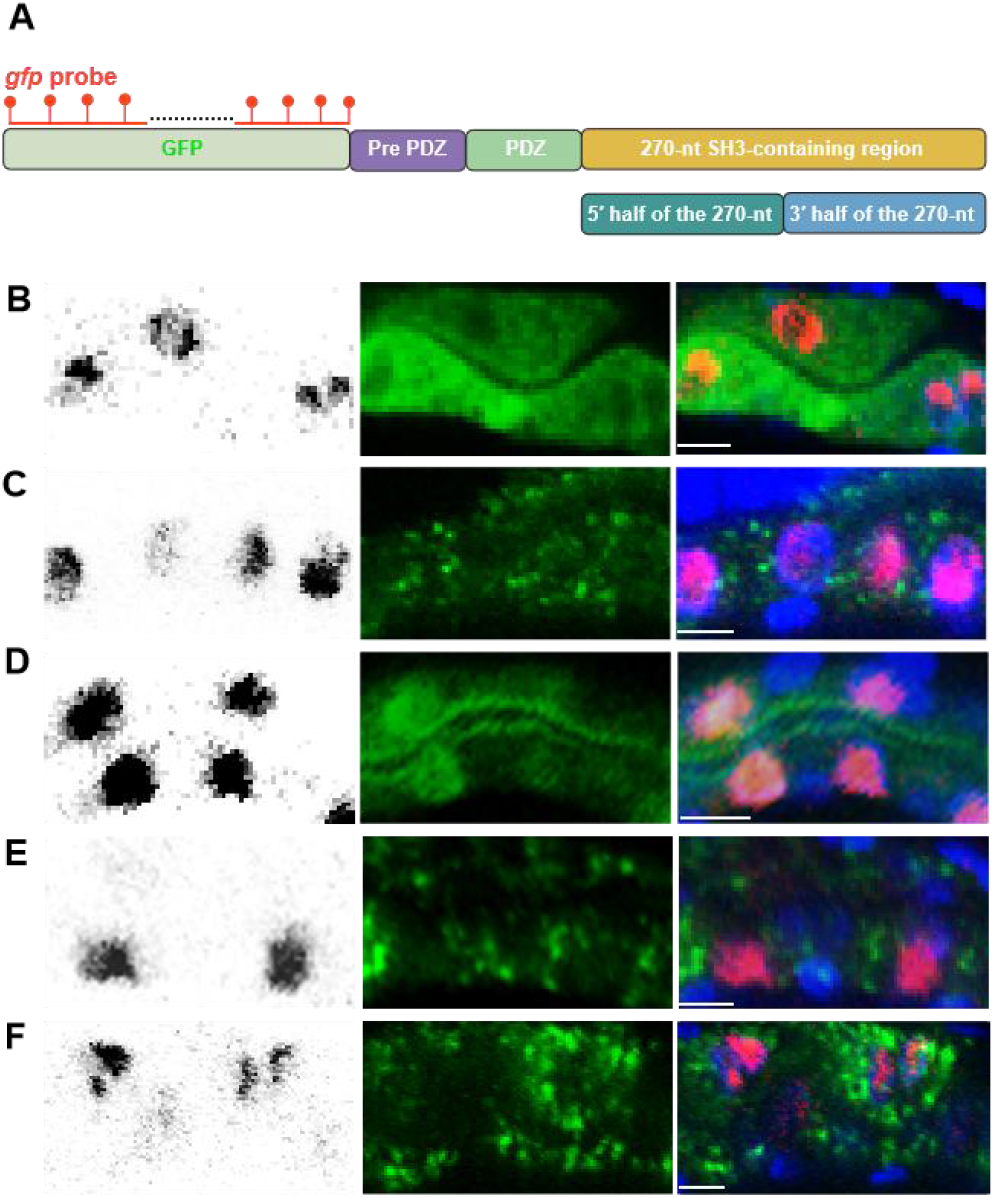
A 270-nt coding sequence within magu-2 exon 3 is required for its apical localization. (A)Schematic of five magu-2 GFP-tagged transgenes. smFISH probes complementary to *gfp* mRNA were used to label transgenic transcripts. (B-F) smFISH probes visualize the distribution of the five magu-2 truncated mRNAs in C. elegans intestinal epithelial cells. Note that the GFP signal was blurred after washing during FISH staining. (B) The exon 3 N-terminal (pre-PDZ) magu-2 truncated transcript displays diffuse cytosolic dispersal. (C) The magu-2 exon 3 transcript encoding the PDZ domain exhibits a uniform cytosolic distribution. (D) The transcript containing the 270-nt region downstream of the PDZ domain, including the SH3 domain, exhibits cytoplasmic distribution and apical enrichment. (E-F) The transcripts corresponding to the 5’ and 3’ halves of the 270-nt region downstream of the PDZ domain show random cytoplasmic distribution. Scale bar: 5 μm for all images

Across all tested *magu-2* exon 3 truncation reporters, smFISH analysis revealed prominent nuclear accumulation of transcripts, likely a potential result of high-level transgenic expression (Fig.2B-F). In the cytoplasm, transcripts encoding the region upstream of PDZ or the PDZ coding sequence exhibited uniform, non-polarized distribution with no discernible apical asymmetry (Fig.2B-C). In contrast, transcripts encoding the 270-nt core fragment displayed asymmetric cytoplasmic distribution, with enrichment at the apical surface of intestinal epithelial cells, in line with the pattern concordant with the apical localization of full-length exon 3 protein observed previously (Fig. 2D) [31]. Consistent with the protein mislocalization phenotypes observed for these truncations, both the 5’- and 3’-135 nt derivatives exhibited a stochastic cytoplasmic distribution, lacking the apical enrichment observed with the full 270-nt fragment (Fig.2E-F).

To explain the loss of apical enrichment and stochastic cytoplasmic distribution upon truncation of the 270-nt core fragment, we performed RNA secondary structure predictions were carried out for the full-length 270-nt fragment and its two 135-nt halves. The independently folded structure of the two 135-nt sub-fragment differed substantially from their conformations within the full-length 270-nt sequence. Meanwhile, the full 270-nt sequence exhibited 5’-3’ sequence complementarity, allowing the formation of stable stem-loop structures that were absent in either sub-fragment alone (Fig.S2A-C). These observations collectively indicate that the 270-nt fragment functions as an integrated localization module, wherein structural integrity is essential for apical targeting.

### 2.3. Live-cell imaging reveals active transport of the 270-nt magu-2 mRNA

To dynamically characterize the intracellular transport dynamics of the 270-nt *magu-2* mRNA, live-cell tracking of the target transcript was performed in transgenic intestinal cells using the PP7/PCP labeling system (Chao et al., 2008; Li et al., 2021). In addition to the prevalent stationary and corralled states typical of cytoplasmic mRNA granules, directed events with net displacements of 1-5 μm were detected among transcripts encoding the 270-nt *magu-2* fragment (Fig. 3A, B, C). According to the criteria for active transport (single linear displacement ≥2 μm, and a velocity range of 0.1-2 μm/s) [32, 33], these findings demonstrate that the 270-nt fragment is sufficient to confer canonical active transport properties on its encoded transcript.

**Figure 3.**
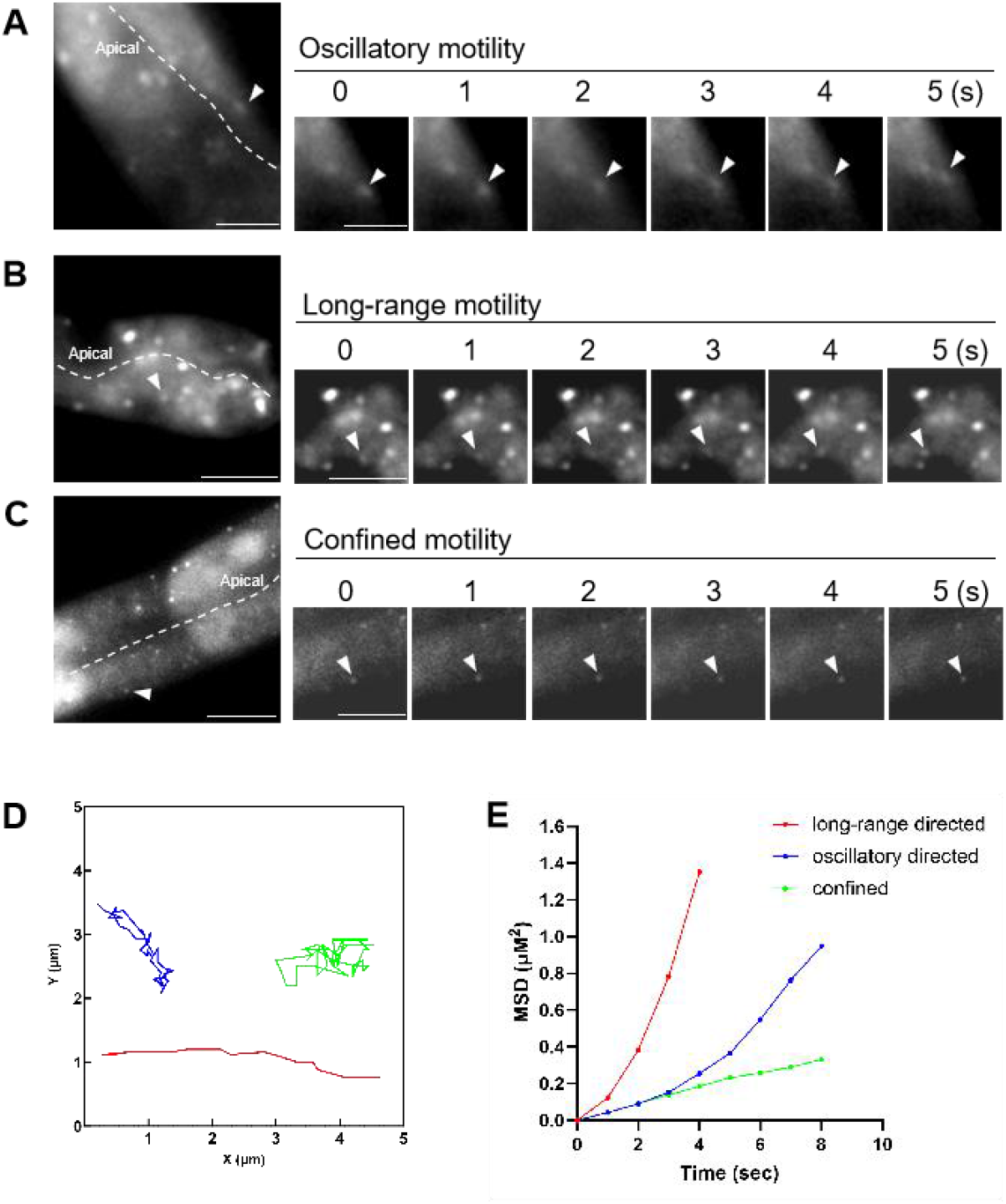
Real-time cytoplasmic dynamics of the 270-nt magu-2 mRNA in C. elegans intestinal cells. (A-C) Representative frames illustrating the three distinct motility patterns of 270-nt magu-2 mRNA particles in intestinal epithelial cells: (A) oscillatory movement, (B) long-range directed transport, and (C) confined diffusion. (D) Two-dimensional tracking traces of 270-nt magu-2 mRNA particles, color-coded over time. (E)Mean squared displacement (MSD) analysis of individual tracked 270-nt magu-2 mRNA particles, stratified by motility phenotype (back-and-forth, long-range directed, confined diffusion).

The primary active transport mode was long-range apical movement, in which individual RNP granules traversed approximately 5 μm of the cytoplasm, moving persistently toward the apical surface (Supplementary Video 1, Fig. 3D). This observation suggests that the 270-nt fragment alone is sufficient to drive long-range directed transport. A second motility pattern, apical-contact oscillation, exhibited periodic bidirectional motility, with each unidirectional trajectory spanning roughly 2 μm, wherein granules initially retreated from the apical periphery toward the mid-cytoplasm and subsequently resumed apical migration, establishing transient contacts with immobile RNP puncta anchored near the apical membrane (Supplementary Video 2, Fig. 3D). In addition, a minor population of RNP granules exhibiting short-range, non-directional restricted motility was occasionally detected in regions of the cytoplasm distant from the apical zone (Supplementary Video 3, Fig. 3D).

Quantitative MSD analysis further resolved these motile behaviors. As illustrated by the representative trajectories, long-range apical runs (red) displayed the steepest slope, characterized by superlinear scaling with a diffusion exponent α of 1.3 (1.2–1.4), indicative of processive, motor-driven transport. Apical-contact oscillations (blue) exhibited a moderately elevated α of approximately 1.1, also consistent with active, non-diffusive motility. Conversely, basal confinement trajectories (green) showed a pronounced sublinear profile with α of 0.6 (0.5–0.7), indicating restricted diffusion within a confined cytoplasmic domain (Fig. 3E). Taken together, these dynamics of the 270-nt-encoded *magu-2* mRNA were in agreement the directed transport phenotypes previously reported for the full-length transcript [31], supporting a model in which the 270-nt mRNA fragment serves as a transport element that engages the conserved mRNA trafficking machinery.

### 2.4. The Dynein/BicD/Egal Axis Mediates Apical Targeting of the 270-nt magu-2 mRNA and MAGU-2 protein

mRNA active transport typically requires the assembly of multiprotein complexes that are propelled by transport factors along the cytoskeleton. The 270-nt fragment within *magu-2* exon 3 constitutes a cis-element sufficient for its apical mRNA localization; however, the trans-acting machinery driving this process remains undefined. While the evolutionarily conserved dynein/BicD/Egl axis has been recognized as a core component of microtubule-dependent mRNA transport [34-37], its involvement in *magu-2* mRNA trafficking, specifically that mediated by the 270-nt element, remained untested. To address this, the localization of both the 270-nt magu-2 transcript and its encoded protein was evaluated following RNAi-mediated knockdown of the C. elegans homologs *dhc-1, bicd*, and *egal*.

In control animals, both the 270-nt-encoded *magu-2* mRNA and MAGU-2 protein exhibited overlapping enrichment at the apical surface of intestinal epithelial cells (Fig. 4A). The most pronounced phenotype was observed following depletion of the RNA-binding protein EGL-1 (*egal* RNAi), which triggered the accumulation of the 270-nt *magu-2* transcript in large cytoplasmic aggregates, whereas the protein adopted a largely diffuse cytosolic distribution interspersed with large puncta representing sites of mRNA-protein co-localization (Fig. 4B). Quantitatively, 96% of *egal* RNAi animals exhibited transcript mis-localization, a frequency higher than that of protein mislocalization, indicating that defects in transcript targeting precede protein mislocalization (Fig. 4F). Similarly, knockdown of the dynactin subunit BICD (*bicd* RNAi) resulted in frequent mislocalization of both the 270-nt magu-2transcript and MAGU-2 protein relative to controls (Fig. 4F), recapitulating transcript aggregation and diffuse protein distribution with small cytosolic puncta showing mRNA-protein colocalization, albeit with a lower penetrance than that observed following *egal* RNAi (Fig. 4C). At the same time, knockdown of the dynein heavy chain DHC-1 (*dhc-1* RNAi) induced defects of comparable severity to that of *bicd* RNAi, with both occurring at markedly higher frequencies than in control animals (Fig. 4F). Following DHC-1 knockdown the 270-nt *magu-2* transcript and MAGU-2 protein predominantly adopted a diffuse cytosolic distribution interspersed with small aggregates (Fig. 4D). Next, to disrupt the microtubule networks in intestinal epithelial cells, animals were treated with the microtubule-depolymerizing agent nocodazole. This treatment triggered frequent mis-localization of both the 270-nt *magu-2* transcript and MAGU-2 protein relative to controls (Fig. 4F). Specifically, the protein was distributed as small, dispersed puncta across the cytoplasm, whereas the transcript accumulated in small aggregates (Fig. 4E). Overall, these results support a role for the dynein/BicD/Egl axis and intact microtubules in mediating apical targeting of the 270-nt *magu-2* transcript and MAGU-2 protein (Fig. 4G). The observation that transcript mislocalization preceded protein mislocalization further implies that mRNA trafficking may prime subsequent apical enrichment of the encoded protein.

**Figure 4.**
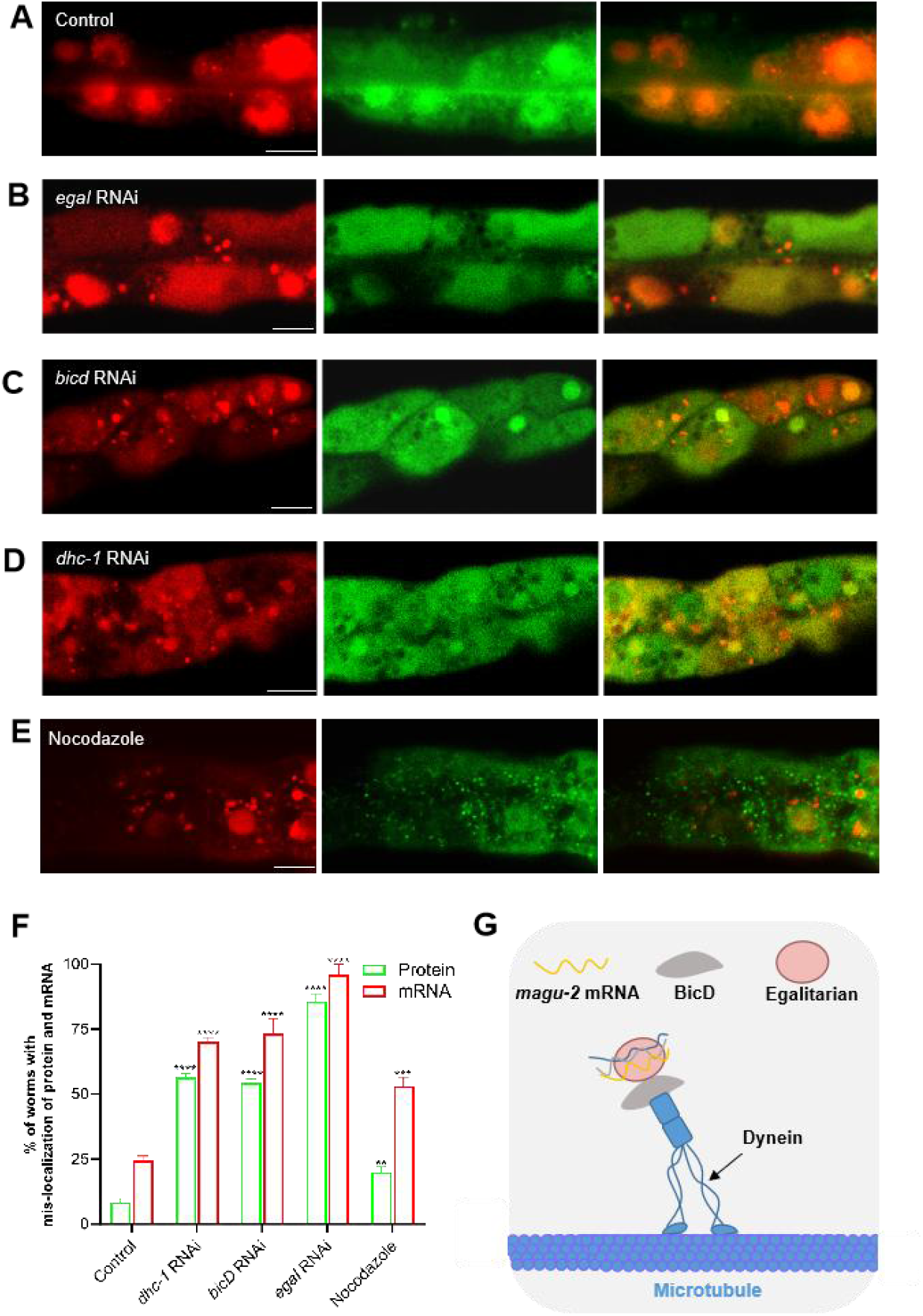
Localization of the 270-nt magu-2 mRNA and protein after perturbation of Dynein/BicD/Egalitarian function or microtubules. (A-E) Confocal images of C. elegans intestinal epithelial cells under control conditions, following RNAi-mediated knockdown of bicd, egal, or dhc-1, or after pharmacological disruption of microtubule networks with nocodazole. The 270-nt truncated magu-2 transcript is labeled with PP7/PCP-Tomato system (red), and the corresponding truncated MAGU-2 protein is tagged with GFP (green). In control animals, both signals exhibit robust co-localization at the apical surface (yellow in merge panels); in all perturbation conditions, apical enrichment is abolished, with both transcripts and protein redistributing to discrete cytoplasmic aggregates and diffuse cytosolic pools. (F)The bar graph indicates the percentage of animals with abnormal protein and mRNA localization. Data represent mean ± SD, n = 3 (10 animals per experiment); **p < 0.01, ***p < 0.001; ****p < 0.0001.

### 2.5. The 270-nt Element Is Dispensable for Epithelial Polarity

To investigate the physiological role of the 270-nt regulatory element within *magu-2* exon 3, a precise deletion allele was generated using CRISPR-Cas9-mediated genome editing, resulting in the *magu-2*(ex3Δ270) mutant strain. This mutant provided a genetic model for assessing the contribution of the conserved exon 3 sequence to epithelial polarity and normal development under standard laboratory conditions.

The subcellular localization of representative polarity markers, including the apical membrane protein PGP-3, the junctional marker AJM-1, and the basolateral marker MRP-1, was first examined in mutant animals (Fig. 5A). As depicted in Fig. 5C–D, all three markers exhibited localization patterns indistinguishable from those observed in wild-type animals, indicating that the overall organization of epithelial domains remained intact in *magu-2*(ex3Δ270) mutants. To further evaluate the potential effects of this deletion on organismal development, brood size was quantified, and no significant difference was noted between *magu-2*(ex3Δ270) mutants and N2 controls (Fig. 5B). These findings conjointly suggest that the 270-nt element is dispensable for maintaining epithelial polarity or supporting normal development under standard culture conditions, further implying that its function may be compensated for by redundant regulatory mechanisms.

**Figure 5.**
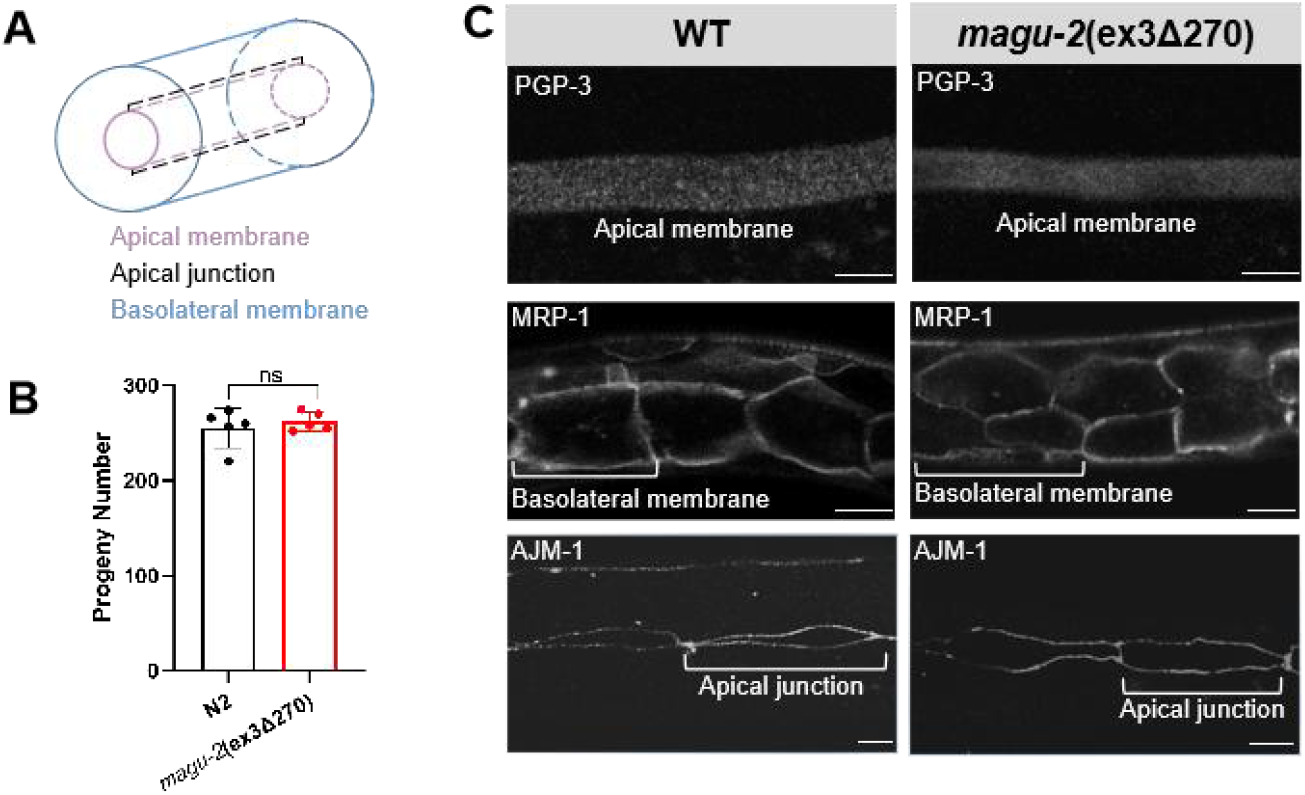
Deletion of the 270-nt element does not disrupt epithelial polarity or brood Size. (A)Schematic of intestinal epithelial cell polarity, illustrating the apical, junctional, and basolateral domains. (B) Quantification of brood size in magu-2(ex3Δ270) mutants compared with wild-type N2 controls. Each data point represents the total progeny count from a single animal; n=5 biologically independent animals per group. Data represent mean ± SD. (C) Images of the basolateral marker MRP-1::GFP, the junctional marker AJM-1::GFP, and the apical marker PGP-1::mCherry in wild-type N2 controls and magu-2(ex3Δ270)mutants. Scare bar: 5 μm for all images.

## 3. Discussion

The establishment and maintenance of epithelial polarity are contingent upon an evolutionarily conserved protein machinery that is fundamentally preserved across metazoans. Although the critical importance of epithelial polarity proteins in establishing and maintaining tissue architecture has been well documented, the contribution of RNA transport to the apical targeting of polarity determinants remains underexplored. MAGU-2 is one such conserved component—the *C. elegans* ortholog of mammalian PALS1 and Drosophila Stardust (Sdt)—and serves as a member of the Crumbs complex [30]. This study provides several new insights into the cis-regulatory determinants governing the coordinated polarization of *magu-2* mRNA and MAGU-2 protein at the apical surface.

Within MAGU-2 exon 3, two distinct sequence segments that can each independently drive apical localization of a GFP reporter were identified; one of these segments encompasses the conserved SH3 domain. To the best of our knowledge, although numerous studies have reported that individual proteins contain multiple apical-targeting determinants, these elements are typically cooperative, enhancing, or otherwise not fully independent. For example, apical localization of the well-studied membrane protein ERM-1 depends on the coordinated action of its FERM domain, which mediates PIP2-dependent membrane association, and C-terminal phosphorylation, which fine-tunes apical dynamics [38]. In contrast, MAGU-2 exon 3 represents an unusual case in which two autonomous, redundant apical-targeting modules coexist within a single exon.

A previous study has evinced that both *magu-2* exon 3 mRNA and the encoded protein are enriched at the apical surface [31]. However, the cis-acting sequence element responsible for mRNA localization remains unknown. An outstanding question is whether the *cis*-acting element responsible for mRNA localization is contained within one of these two sequence segments, or whether it is separate from the sequence elements that direct protein targeting. It is worthwhile emphasizing that smFISH analysis showed that the mRNA corresponding to the 270-nt fragment encompassing the SH3 domain was enriched at the apical surface, suggesting that the cis-acting element responsible for *magu-2* exon 3 mRNA localization resides within this region. Further truncation resulted in diffuse cytoplasmic distribution of both the mRNA and the encoded protein, indicating that the shorter fragment is insufficient to support proper subcellular localization and that the sequence information required for localization is distributed within across the 270-nt segment. Generally, mRNA localization elements are located in the 3’UTR or 5’UTR, although coding-sequence localization signals have also been reported [39]. The results of this study suggest that the cis-acting element responsible for *magu-2* exon 3 mRNA apical localization resides within the coding region and overlaps with the sequence encoding the SH3 domain, indicating a non-canonical localization mechanism. This region may therefore integrate information regulating both protein targeting and mRNA localization, potentially through RNA sequence motifs, local secondary structures, or interactions with RNA-binding and translation-associated factors.

This raises the question of how this mRNA achieves its subcellular localization. To address this, its transport behavior was examined from the perspective of RNA trafficking. The observed unidirectional movements over 2-5 μm, together with the MSD analysis, were consistent with motor-driven intracellular trafficking. These findings suggest implies that the cis-acting sequence element identified within the 270-nt sequence may recruit specific RNA-binding proteins and cytoskeletal transport machinery to direct mRNA movement to the apical region. Given that this element overlaps with the conserved SH3-encoding sequence, the *magu-2* transcript may represent a non-canonical example of coding-sequence-mediated mRNA localization that couples subcellular targeting with protein-coding information.

We propose that the subcellular localization of *magu-2* mRNA is mediated by a cis-acting localization element that recruits specific RNA-binding proteins, which in turn link the transcript to the cytoskeletal transport machinery, thereby enabling its directed transport. In *Drosophila*, the subcellular localization of *crb* mRNA has been shown to depend on recognition by EgI, the Drosophila ortholog of EGAL-1, through its localization element (LE), while the localization of *sdt* mRNA has also been reported to rely on dynein-mediated transport [40]. However, whether a similar mechanism operates in *magu-2* mRNA localization has not yet been investigated. To identify transport factors involved in the trafficking of *magu-2* mRNA, the roles of DHC-1, BICD, and EGAL-1 were examined, given that their *Drosophila* orthologs have previously been implicated in polarized mRNA localization [34, 41]. Additionally, microtubule integrity was disrupted to assess their contribution to *magu-2* mRNA and protein localization. Among these perturbations, *egal-1* RNAi generated the most pronounced phenotype. Assuming comparable RNAi efficiency, the similar severity observed following *dhc-1* and *bicd* knockdown suggests that these factors may act within the same microtubule-dependent transport pathway but at different steps. In particular, the stronger phenotype caused by EGAL-1 depletion may indicate that it functions at a more upstream or rate-limiting step in this pathway. Notably, the penetrance of the mRNA phenotype was consistently higher than that of the protein phenotype across these RNAi conditions, suggesting that *magu-2* mRNA is more sensitive to disruption of these transport factors. This finding implies that DHC-1, BICD, and EGAL-1 may directly participate in mRNA recognition and transport, whereas the protein localization defect may be influenced to a greater extent by downstream processes and therefore appears less severe. Besides, under these conditions, the 270-nt *magu-2* mRNA predominantly accumulated in aggregates, whereas the corresponding protein exhibited a diffuse distribution, with both losing their normal apical enrichment. This divergence between RNA and protein localization suggests that mRNA trafficking, local translation, and subsequent protein anchoring may not be fully coupled, but are instead are regulated by distinct steps within the same transport framework. Taken together, these findings support a model in which DHC-1, BICD, and EGAL-1 cooperate to mediate the apical transport of *magu-2* mRNA in a microtubule-dependent manner, highlighting the importance of an intact transport machinery for proper subcellular localization.

CRISPR-mediated deletion of the 270-nt magu-2 sequence did not affect apical-basal polarity or worm development, indicating that this region is functionally redundant. This redundancy is further supported by the presence of an additional apical-targeting element that may partially compensates for the loss of this 270-nt region at the protein localization level. Importantly, previous studies have demonstrated that MAGU-2 is dispensable for the establishment and maintenance of apical-basal polarity in the *C. elegans* intestine and is not essential for worm development (Castiglioni et al., 2022). Interestingly, overexpression of the fragment of *magu-2* containing PDZ and SH3 domain-containing fragment causes defects in intestinal tubulogenesis, suggesting that this region may mediate interactions with specific binding partners and that its overexpression may sequester these factors in a dominant-negative manner.

Although MAGU-2 is not essential for *C. elegans*, the identification of an mRNA localization element within its SH3 domain, together with the conservation of this region and the cross-species conservation of MAGU-2 localization and protein-protein interactions, highlights the biological relevance of this study for understanding the mechanisms underlying epithelial polarity.

## 4. MATERIALS AND METHODS

### 4.1. C. elegans strains

*C. elegans* culture, maintenance, and genetic crosses were carried out as described [42]. Strains were routinely maintained at 20-22 °C. All transgenic and mutant strains used in this study were derived from the N2 background, and the complete list of strains is provided in Table S1.

### 4.2. Construction of transgenic and mutant strains

For molecular cloning, the PDZ-SH3 fragment (*magu-2* exon 3, nucleotides 355–828), *magu-2* exon 3 [1–354], *magu-2* exon 3 [829–840], *magu-2* exon 3 [355–558] and *magu-2* exon 3 [559–828] were generated using the *Pact-5::GFP::magu-2 (exon 3)::PP7* plasmid as the template. The target fragments and vector backbone were separately amplified and assembled by seamless cloning.

To generate the truncated constructs, the *Pact-5::GFP::magu-2 (exon 3)::PP7* plasmid was used as the template to generate *magu-2* exon 3 del[355–828]. The *Pact-5::GFP::magu-2 (exon 3 559–828)::PP7* plasmid was used as the template to generate *magu-2* exon 3 [559–693] and *magu-2* exon 3 [694–828]. All constructs were generated by overlap PCR using Vazyme high-fidelity DNA polymerase. Briefly, primers were designed with 10–15 bp overlapping complementary sequences at their 5′ ends. The resulting PCR products were treated with DpnI and then transformed into competent cells.The primer sequences are listed in Table S2.

All constructs were sent to SunyBiotech for microinjection. The *magu-2* exon 3 [559–828] (Syb11566) knockout mutant strain was generated by SunyBiotech.

### 4.3. smFISH procedure

RNA probes against GFP were designed using the Stellaris RNA FISH Probe Designer (https://www.biosearchtech.com/support/tools/design-software/stellaris-probe-designer) and custom-synthesized by Biosearch Technologies, with conjugation to CAL Fluor Red 610 dye. Single-molecule fluorescence in situ hybridization (smFISH) was performed according to standard protocols with minor modifications. Briefly, probes were dissolved in RNase-free TE buffer to a final stock concentration of 250 μ M. Synchronized L1 larvae were cultured on OP50 bacteria until 24 h after the mid-L4 stage and then collected using M9 buffer. Afterward, worms were fixed in 3.7% formaldehyde in 1×PBS containing 0.1% Tween-20 for 15-45 min at room temperature, permeabilized with 0.1% Triton X-100, and dehydrated in 70% ethanol overnight at 4 °C. After prehybridization in wash buffer, samples were hybridized with 0.1-0.5 μ M probes in hybridization buffer overnight at 37 °C in the dark. Following hybridization, samples were washed, counterstained with DAPI, and mounted in Antifade Prolong Gold medium for imaging.

### 4.4. RNAi knock down and drug treatment experiments

To generate RNAi constructs targeting *dhc-1, bicd*, and *egal-1*, the target fragments were amplified from *C. elegans* cDNA by PCR and inserted into the L4440 vector linearized with NotI and XbaI using seamless cloning. The primer sequences are listed in Table S2. RNAi plates were prepared by spreading RNAi clone cultures onto NGM plates supplemented with 2 mM IPTG and 50 μg/mL carbenicillin. Ten L4-stage worms were transferred onto the RNAi plates and incubated at 22°C for 2 days. The F1 progeny at the L2–L3 stage were subsequently selected for phenotypic analysis. Worms fed with the empty L4440 RNAi vector served as the negative control.

For drug treatment, L3-stage worms were treated in M9 buffer supplemented with 60 μg/mL nocodazole and a small amount of OP50 for 2 h. The worms were then centrifuged at 5000 rpm for 5 min, following which the supernatant was discarded, and the pellets were resuspended in fresh M9 buffer and allowed to recover for 1 h before being mounted and observed under a confocal microscope.

### 4.5. Fluorescence microscopy

For live-cell and smFISH imaging, worms were anesthetized in M9 buffer containing 5 mM levamisole and mounted on glass slides. Imaging was performed using a Sunny IRX60 confocal fluorescence microscope equipped with a 60× oil-immersion objective. GFP, Tomato/Cy3, and DAPI were excited with 488 nm, 561 nm, and 405 nm lasers, respectively.

For time-lapse imaging, live worms were mounted on 2% agarose pads and anesthetized with 0.1% tetramisole. The coverslip was sealed with Vaseline. Images were acquired using a Leica MICA microscope in widefield mode with a 60× water-immersion objective, at intervals of 1 s.

### 4.6. Single-particle tracking analysis

Time-lapse images were analyzed using Fiji. Positional data for individual RNA particles were obtained using MTrackJ. Based on the positional data of each RNA particle, mean square displacement (MSD) was calculated using MATLAB. A custom-written MATLAB script was used for the analysis (Park et al., 2014).

### 4.7. Statistical analysis

Statistical analyses were performed using GraphPad Prism 10. Data presented in bar graphs are expressed as mean ± SEM. Each group represents the average of at least three independent experiments. P values were calculated using Student’s t-test, and statistical significance was defined as follows: *p < 0.05, **p < 0.01, ***p < 0.001, and ****p < 0.0001.

## 5. Conclusions

In summary, our study demonstrated that a 270-nt coding-sequence element within *magu-2* exon 3, encompassing the conserved SH3 domain, functions as a cis-acting localization module that directs the apical targeting of both *magu-2* mRNA and its encoded MAGU-2 protein. Moreover, Motor-driven trafficking analysis and RNAi perturbation reveal that DHC-1, BICD, and EGAL-1 cooperate in a microtubule-dependent pathway to mediate apical mRNA transport. The dissociation between RNA aggregate formation and diffuse protein mislocalization under transport disruption implies that mRNA trafficking, local translation, and protein anchoring are partially uncoupled yet integrated processes. Although CRISPR deletion of this region yields no polarity or developmental defect due to functional redundancy, the conservation of the SH3-adjacent localization signal across metazoans underscores its evolutionary significance. These findings expand the scope of coding-region-mediated RNA targeting and provide a framework for understanding how epithelial polarity structural proteins achieve subcellular precision beyond canonical UTR-based mechanisms.

## Supplementary Materials

The following supporting information can be seen in the attached supplementary file.

## Author Contributions

**Z.L. designed the study. Z.L., Q.J., Z.W., and Y.S. carried out the experiments. Z.L., Z.W., and Y.S. analyzed the data. Z.L. and Q.J. drafted and revised the manuscript**.

