## Supplemental Data 1 for "A Cis-Element in *magu-2* Mediates Apical mRNA Localization via Active Transport in *C. elegans*"

**Table S1 Strain list**

| Strain name | Genotype | Reference |
|---|---|---|
| N2 | wild-type | Brenner, 1974 (CGC) |
| HJZ051 | *mauls03[Pelt-2::NLS::tdPCP::tdTomato,Prol-6::rol-6(su1006)]* | Li et al., 2021 |
| HJZ070 | *mauEX28[Pact-5::GFP::magu-2 Exon3::PP7,Prol-6::rol-6(su1006)]* | Li et al., 2021 |
| OG472 | *drSi2 [vha-6p::3XFLAG::pgp-3/CFTR(wt)::mCherry::unc-54 3'UTR] II* | He L, et al.,2012 (CGC) |
| AY173 | *acEx173 [mrp-1::GFP]* | Lalsiamthara J, et al.,2022 (CGC) |
| PS3729 | *syIs78 [ajm-1::GFP + unc-119(+)]* | CGC |
| LIZ01 | *EX1[Pact-5::GFP::magu-2 Exon3(355-828)::PP7,Prol-6::rol-6(su1006)]* | This study (SunyBiotech) |
| LIZ02 | *EX2[Pact-5::GFP::magu-2 Exon3(Δ355-828)::PP7,Prol-6::rol-6(su1006)]* | This study (SunyBiotech) |
| LIZ03 | *EX3[Pact-5::GFP::magu-2 Exon3(1-354)::PP7,Prol-6::rol-6(su1006)]* | This study (SunyBiotech) |
| LIZ04 | *EX4[Pact-5::GFP::magu-2 Exon3(829-840)::PP7,Prol-6::rol-6(su1006)]* | This study (SunyBiotech) |
| LIZ05 | *EX5[Pact-5::GFP::magu-2 Exon3(355-558)::PP7,Prol-6::rol-6(su1006)]* | This study (SunyBiotech) |
| LIZ06 | *EX6[Pact-5::GFP::magu-2 Exon3(559-828)::PP7,Prol-6::rol-6(su1006)]* | This study (SunyBiotech) |
| LIZ07 | *EX7[Pact-5::GFP::magu-2 Exon3(559-693)::PP7,Prol-6::rol-6(su1006)]* | This study (SunyBiotech) |
| LIZ08 | *EX8[Pact-5::GFP::magu-2 Exon3(694-828)::PP7,Prol-6::rol-6(su1006)]* | This study (SunyBiotech) |
| LIZ09 | *mauls03[Pelt-2::NLS::tdPCP::tdTomato,Prol-6::rol-6(su1006)]; EX6[Pact-5::GFP::magu-2 Exon3(559-828)::PP7,Prol-6::rol-6(su1006)]* | This study |
| LIZ10 | *Syb11566[magu-2(exon3Δ559–828)] II* | This study (SunyBiotech) |
| LIZ11 | *Syb11566[magu-2(exon3Δ559–828)] II;drSi2 [vha-6p::3XFLAG::pgp-3/CFTR(wt)::mCherry::unc-54 3'UTR] II* | This study |
| LIZ12 | *Syb11566[magu-2(exon3Δ559–828)] II; acEx173 [mrp-1::GFP]* | This study |
| LIZ13 | *Syb11566[magu-2(exon3Δ559–828)] II; syIs78 [ajm-1::GFP + unc-119(+)]* | This study |

**TableS2 Primer list**

| Primer name | Primer sequence(5’-3’) |
|---|---|
| magu2-exon3-2-F | CTTGGCGCAACGGTTCGAA |
| magu2-exon3-2-R | CGAGGAACTAGGAATTAATCCTGCC |
| magu2-exon3-3-F | AAAGAAGAACAAGACACCGAAC |
| magu2-exon3-4-R | TGGTGCAACTACAAAAGTAATTGG |
| magu2-exon3-5-F | GAGGATAGTTATTTTAGACAACAataaGGACAGCG |
| magu2-exon3-5-R | TTGTTGTCTAAAATAACTATCCTCATCCTTCACAAC |
| magu2-exon3-6-F | cAGCTTGACGAAGTAATTATATCTGTTG |
| magu2-exon3-6-R | ATAACTATCCTCATCCTTCACAAC |
| magu2-exon3-7-F | GAGTTGGCAATGAAATTTCAAAG |
| magu2-exon3-7-R | TTGTTGTCTAAACGAGGAACT |
| Vector-2-F | GGATTAATTCCTAGTTCCTCGGGACAGCGTGCTAAtttgtaatG |
| Vector-2-R | CGAACCGTTGCGCCAAGTTTGTATAGTTCgTCCATGCCA |
| Vector-3-R | CGGTGTCTTGTTCTTCTTTTTTGTATAGTTCgTCCATGCCA |
| Vector-4-F | TTACTTTTGTAGTTGCACCAGGACAGCGTGCTAAtttgtaatG |
| Vector-6-F | GAAGGATGAGGATAGTTATGGACAGCGTGCTAAtttgtaatG |
| Vector-6-R | GATATAATTACTTCGTCAAGCTGTTTGTATAGTTCgTCCATGCC |
| Vector-7-F | GTTCCTCGTTTAGACAACAAGGACAGCGTGCTAAtttgt |
| Vector-7-F | GAAATTTCATTGCCAACTCTTTGTATAGTTCgTCCATGCC |
| FP-SH3-truncate-1 | CCTTGTAAAGAGGGACAGCGTGCTAAtttgtaatG |
| RP-SH3-truncate-1 | TAGCACGCTGTCCCTCTTTACAAGGTACATACACG |
| FP-SH3-truncate-2 | cGAACTATACAAAGAGTTGGCAATGAAATTTCAAAG |
| RP-SH3-truncate-2 | TGCCAACTCTTTGTATAGTTCgTCCATGCCATG |
| dhc-1-RNAi-F | GCGGCCGCTCTAGAAAAGGAAGGAGCTCAACGACA |
| dhc-1-RNAi-R | CCGGTGGATCCACTAGCCTTTCCTTCCTGGGTCTTC |
| bicd-1-RNAi-F | GCGGCCGCTCTAGAATGCTCGTCAACAAAAAGACG |
| bicd-1-RNAi-R | CCGGTGGATCCACTAGTTGCAGTGAACATTGCCCTA |
| egal-1-RNAi-F | GCGGCCGCTCTAGAATCATTTCCAGGGTCATCCAT |
| egal-1-RNAi-R | CCGGTGGATCCACTAGTTTCTCCATCAACTCCGACC |

**Figure S1. Colocalization analysis of MAGU-2 truncation constructs with the apical marker PGP-3.**

**(A)GFP fused to the 270-nt SH3-encoding region downstream of the PDZ domain (green) and PGP-3 (red). The merged image reveals co-localization at the apical surface.**

**(B)GFP fused to the N-terminal region of exon 3 upstream of the PDZ domain (green) co-localizes with PGP-3 (red) at the apical surface, as indicated by the yellow signal in the merged image. Scare bar: 5 μm for all images.**

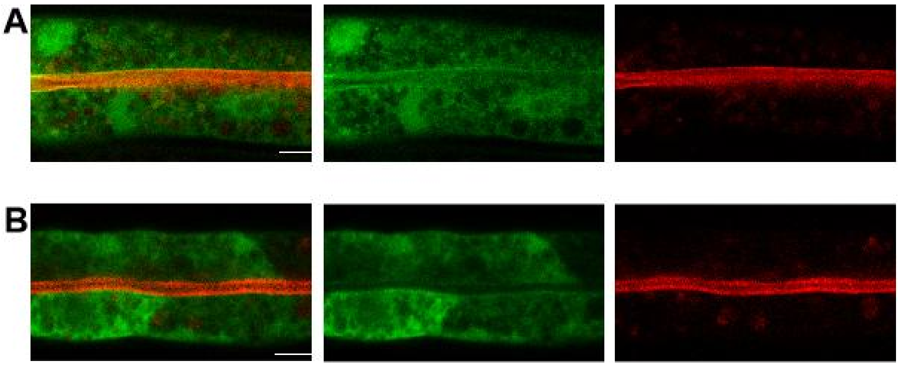

**Figure S2. mfold-predicted RNA secondary structure**

**(A)Predicted RNA secondary structure of the 270-nt SH3-encoding region downstream of the PDZ domain.**

**(B-C) Predicted RNA secondary structure of the 5'and 3' halves of the 270-nt region downstream of PDZ.**

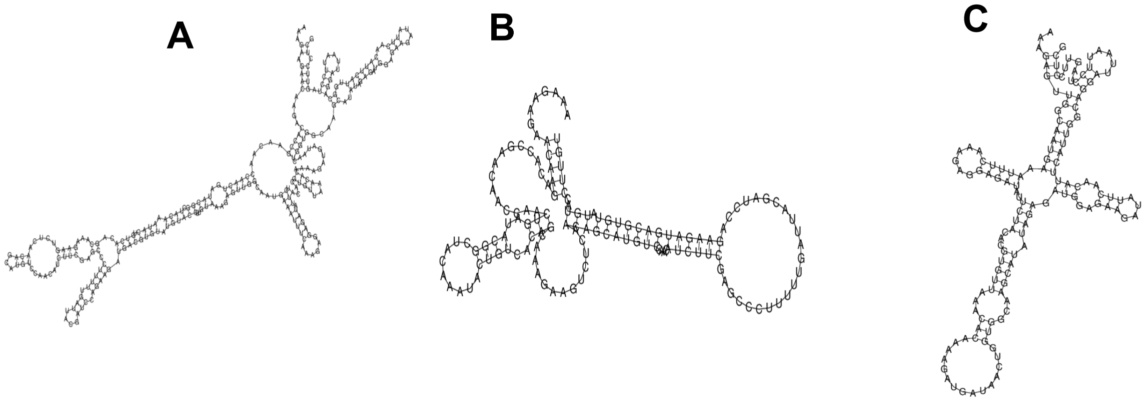
